## Supplemental Material for "Regulation of Ebola GP conformation and membrane binding by the chemical environment of the late endosome"

### Supporting Information

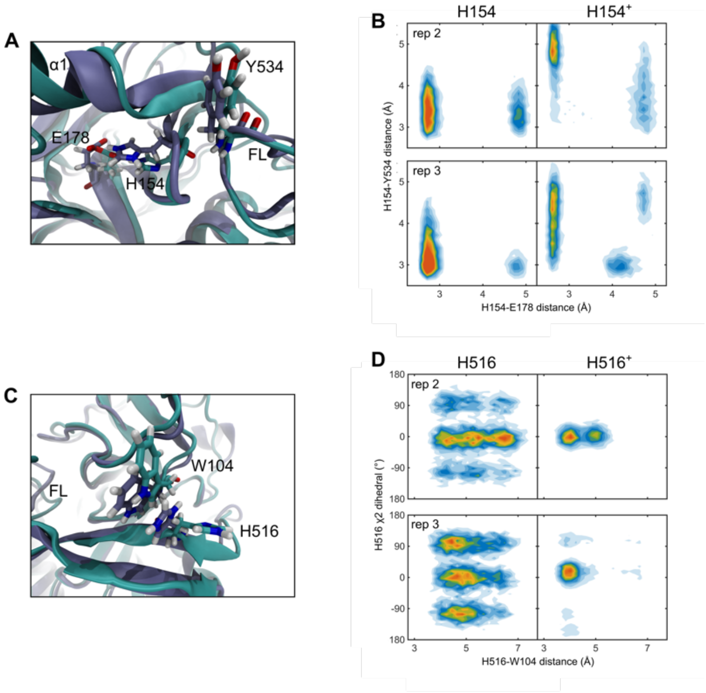


**Figure S1. Replicate data from MD simulation.** (**A**) Zoomed-in view of (purple) protonated and (cyan) deprotonated simulation frames depicting an overlay of the interactions between H154, E178, and Y534. (**B**) As in **Figure 4D**, contour plot indicates the position of H154 in terms of the distance between the H154 and E178 sidechains, and between the H154 and Y534 backbones. Data are shown for replicas two and three of the protonated ($H154^+$) and deprotonated (H154) simulations. (**C**) Zoomed-in view of (purple) protonated and (cyan) deprotonated simulation frames depicting an overlay of the interactions between H516 and W104. (**D**) As in **Figure 4G**, contour plot indicating the orientation of H516 in terms of the $\chi_2$ sidechain dihedral and the distance between the H516 and W104 side chains. Data are shown for replicas two and three of the protonated ($H154^+$) and deprotonated (H154) simulations.

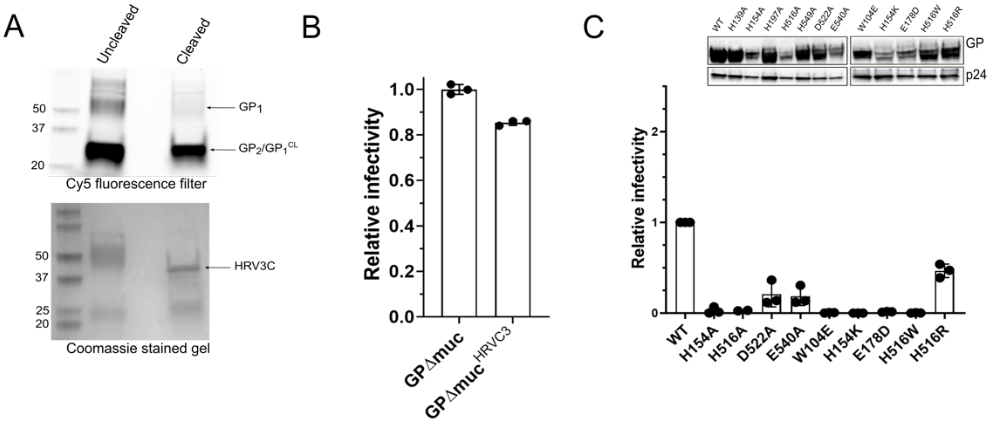


**Figure S2. Proteolysis, labelling, and infectivity of GP variants.** (**A**) Denaturing polyacrylamide gel showing uncleaved GPΔTM and $GP^{CL}$ (cleaved with HRV3C) imaged under Cy5 fluorescence (top) and stained with Coomassie (bottom). (**B**) Comparison of relative infectivity of GPΔmuc, and GPΔmuc with the HRV3C cleavage site. (**C**) (top) Western blot of pseudovirions showing expression of GPΔmuc (with the HRV3C cleavage site) and p24. (bottom) Relative infectivity of lentiviral pseudoparticles containing the mutant GPΔmuc. All infectivity experiments were performed with three biological replicates, each measured in triplicate. Data points represent the mean infectivity determined across biological replicates.

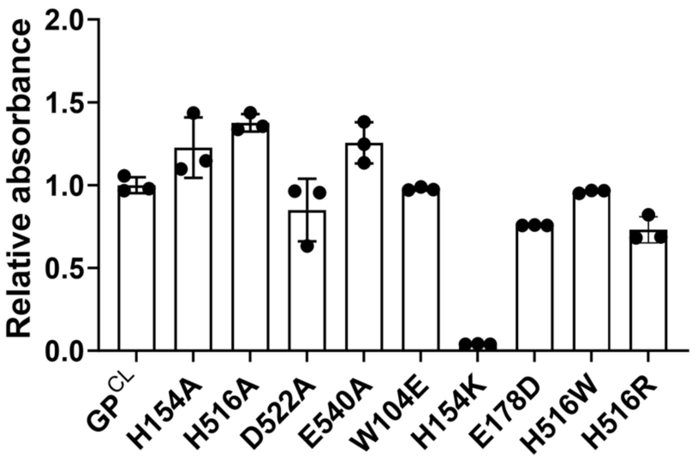


**Figure S3. Antigenicity of mutant GPs.** (**A**) ELISA of wild-type and mutant GPΔTM proteins with mAb KZ52. Except for H154K, introduction of mutations in GPΔTM does not have a significant impact on the antigenicity of GP. ELISAs were carried out in triplicate. Data points represent absorbance values at 450 nm. Bars and error bars represent the mean and standard deviations.

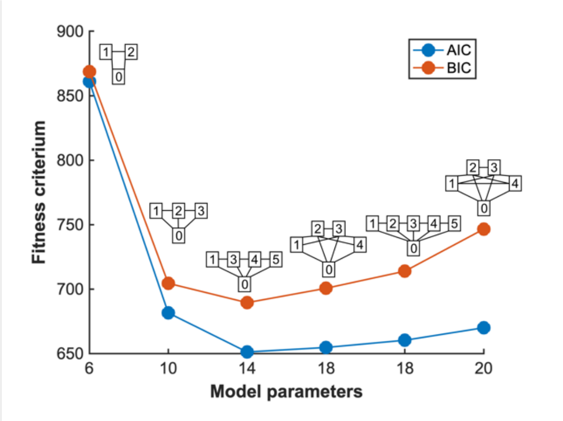


**Figure S4. Model selection through minimization of the AIC and BIC.** smFRET trajectories from wild-type $GP^{CL}$ were fit to a series of different models by maximum likelihood optimization using the MPL algorithm (see Materials and Methods). The maximized likelihood estimated for each model was corrected for differing numbers of model parameters using the AIC and BIC criteria. Under this procedure, the model with best fitness among those considered will generate minimum AIC and BIC values. Both criteria identified the 5-state linear model as providing the best representation of the data among the models considered. Overlaid on the plot are schematic representations of the kinetics models considered. In all models, the 0 state corresponds to the 0-FRET state, with the others reflecting non-zero FRET states. Lines represent connections (allowed transitions) between states. All non-zero FRET states are connected to the 0-FRET state since photobleaching is seen from all FRET states. TDPs shown in **Figures 2** and **5-7** support the identification of a linear model.
